# Using Monoids for the Integration of Genomic and Metabolic Parameters in the Prediction of Phenotypes in Regulatory Cascades

**DOI:** 10.1101/2025.04.03.646970

**Authors:** Marco Villalba, Yair Romero

## Abstract

Molecular network modeling requires the use of mathematical and computational formalisms for a robust and accurate prediction of phenotypes. Furthermore, there is a need to extend these formalisms to be applied to large-scale molecular networks, thus helping in the understanding of biological complexity. In this work, we propose an extension to the modeling framework known as design principles, developed by M.A. Savageau, which is based on the Power Law Modeling. While valuable to understand several properties of molecular networks, the power law modeling cannot be used to infer kinetic orders, which are typically related to the number of binding sites present in molecules. We modified the traditional approach by incorporating the use of monoids generating a new methodological approach that we call Genotype Arithmetic. This approach solves local geometry, solving networks through monoids and varieties, reducing the computational complexity. The resulting combinatorial object contains a family of geometric points, whose fixed points in the exponent space define a set of constraints determining the correct kinetic order to be used in the power law modeling. This characteristic constitutes a prediction of the binding strength and/or the number of DNA binding sites in regulatory sequences, as well as the reaction orders in enzymatic kinetics. To show the applicability of the present approach, we show how the number of binding sites can be approximated in metabolic pathways formed by 3 to 9 reactions, in allosteric systems of end-product inhibition with intermediate catalytic reactions and one gene inhibition.

**Author summary:** In this work, was revised the Power Law Modeling to analyze and predict kinetic and genomic parameters in molecular networks. It was developed a new approach, called Genotype Arithmetic, which takes advantage from the properties of Monoids found within this formalism. Using this algebraic and geometric technique, were predicted the number of binding sites for various cases associated with the pathway length in the gene allosteric system for end-product inhibition.

## Introduction

A key objective in molecular network modeling is to systematically characterize all genetic and metabolic interactions within a living cell [1], [2], [3], [4], [5], [6], [7], [8], [9], [10], [11], [12], [13], [14], [15], [16]. Genomics has advanced rapidly due to massive genome sequencing technologies and post-genomic methods, which enable the identification of whole-genome properties from a single experiment. These include transcription factor binding sites (TFBSs) identified via ChIP-seq and whole-genome expression analysis through RNA-seq, applied to bulk cellular extracts and, more recently, to single-cell analysis (Single Cell-seq) [17–21]. These advances have accelerated the reconstruction of molecular pathways and networks. Consequently, mathematical and computational formalisms capable of approximating these parameters are highly valuable.

In this work, we build upon the power-law model and incorporate the theory of monoids. The power law model for biological networks, extensively studied and developed by M.A. Savageau, has numerous applications [22–33]. Several approaches aim to infer, or at least narrow down, the range of possible parameter values within a given modeling framework. For example, the power law modeling allows for the prediction of kinetic parameters, such as affinity constants and the production and degradation rates of metabolites and proteins, particularly in cases where the underlying metabolic reactions are described by a system of nonlinear differential equations. However, despite its utility, the power law modeling cannot determine the kinetic orders, which often correspond to the number of binding sites.

Power-law polynomials possess structural properties that offer significant mathematical advantages, particularly their combinatorial properties. In this work, we study a geometric object called a Monoid [34], [35], [36], [37], [38], [39] which has been widely investigated in combinatorial algebraic geometry [34], [35]. This geometric object contains a set of generators that can be computed in polynomial time [40], significantly reducing the computational complexity of solving for kinetic orders in molecular networks. Specifically, the monoid contains a set of points whose coordinates define constraints that can predict binding site values.

Our approach enables the exploration of this structure through a mapping in which one set of values corresponds to the number of binding sites, while another set represents molar concentrations. The monoid structure also includes a set of geometrical bases. The combinatorial properties of these geometries could potentially elucidate new insights into gene regulatory sequences and genomic techniques. For instance, this approach could inspire new alignment methods or the prediction of novel regulatory binding sites [35].

## Materials and methods

### Power Law Modeling

There is a significant opportunity to model cellular networks using the power-law formalism, as shown below in Equation (1). Here, 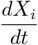 corresponds to the rate law (*mol/s*) for any specific sustainable or stable molecular phenotypes *X*_*i*_’s. The production of this phenotype is governed by the constant rate *α*_*i*_ and species *X*_*j*_ with number of binding sites *g*_*i*_’s, while its degradation is governed by the constant rate *β*_*i*_, and kinetic order *h*_*i*_, see [23–30]. The equation is as follows:

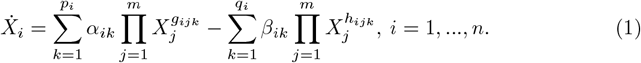

The term 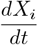 represents the rate of change of molar concentration for any i-th biomolecule in the network. These concentrations belong to formal ring polynomials of Laurent series, denoted by 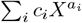. The set of prime polynomials corresponds to an algebraic variety denoted by *Spec*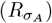, [34–38]. The constants *α*_*ik*_’s and *β*_*ik*_’s are kinetic parameters representing production and degradation rates, respectively, and they also are related to regulatory capacity in the context of TF-DNA interactions (see [21–33]).

Suppose the equation is a rational function. In this case, we rewrite the equation with all exponents in the numerator, either as positive or negative values (Step 2). The exponents *g*_*ijk*_’s and *h*_*ijk*_’s are the kinetic orders, corresponding to the number of binding sites in the regulatory sequences of a transcriptional unity in this formalism. We explicitly quote to M.A. Savageau: “ In general, the exponents in the power laws that characterize classical chemical kinetics are small integer values, as are the exponents in the rational functions that characterize classical biochemical kinetics. In the case of an enzymatic reaction, the largest exponent in the rate law is equal to the number of reactant binding sites on the enzyme (Wyman, 1964) [41], and this is typically equal to the number of subunits in a multimeric protein (Monod et al., 1965). In the case of a regulator that is a multimeric DNA binding protein, the largest exponent is equal to the number of subunits in the regulator molecule multiplied by the number of specific sites on the DNA to which it binds. “ See [33]. Collectively, these variables define a genetic and metabolic network.

### Monoids

The equation is rewritten with all exponents in the numerator, either positive or negative (Step 2 in the workflow). The exponents *g*_*ijk*_’s and *h*_*ijk*_’s represent kinetic orders, corresponding to the number of binding sites for TF-DNA interactions. Additionally, they describe biochemical reaction orders for protein-metabolite and protein-protein interactions. These kinetic orders form a lattice vector (*g*_*ijk*_ and *h*_*ijk*_). From this representation, we construct a Monoid as follows. First, for our methodology, the following formal definitions are necessary:

#### Definition (Lattice-Cone *σ*)

It is the linear space defined by all combinations for the vectors of support space 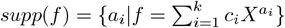 (exponent-space), including addition, subtraction, and scalar multiplication of vectors, if the vectors consist only of integer entries, is called a lattice-cone, we denote it as *σ*, see [34–38].

#### Definition (Dual cone and orthogonal cone)

*σ*^*∨*^ = { v ∈ ℤ^*n*^ :*< v, σ >* ≥ 0 }, see details in [34–38].

#### Definition (Monoid)

Now we intersect the cone with the ndimensional integer space, we write, *σ*^*∨*^ ∩ ℤ^*n*^, we call to this set as a Monoid or as a set *M* of rank *n* or isomorphic to ℤ^*n*^, see [34–38].

#### Lemma (Gordan’s Lemma)

Hilbert basis. Given a cone *σ* and we take the intersection with the *n* dimensional integer space *σ*^*∨*^ ∩ ℤ^*n*^, then this a Monoid, has a finite set of generators, this set of generators is called, the Hilbert basis, see [37, 39].

### Gene Matrix and Fix Points on torus

We construct a Gene Interaction Matrix of lattice vectors Con(*X*_*i*_),, forming vectors for each monomial term in lexicographic order (see [34–38]). The first and last rows of the matrix correspond to the original vectors (Step 3). Using the Boltzmann distribution, we then build a set of orthogonal matrices for Con(*X*_*i*_) (step 4).

These bases define the generators for a Monoid (generator space for binding sites in genome sequences). In this space, we compute fix points with the formula (2), see [34], called fix points on the torus.

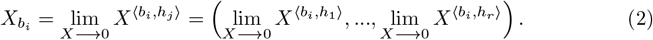

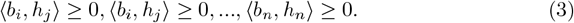

These mappings indicate that the molar concentration of a quantitative phenotype 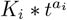 can be represented as the action of an algebraic torus. In other words, molar concentrations belong to orbits on a torus (with t as a complex variable) by the scalar inner product in the exponent space. These facts are justified by the homomorphism of algebras between algebraic and geometric objects, as illustrated in Fig. 2.

**Fig 1.**
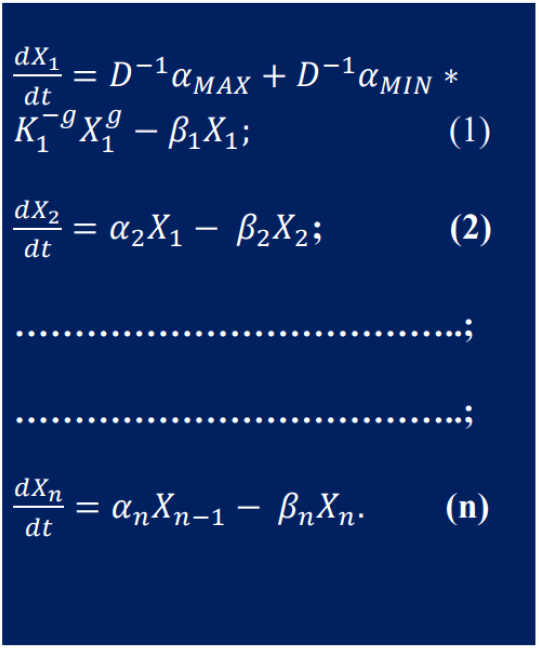
The equation in the figure corresponds to the network scheme for End Product Inhibition for n reactions, the kinetic parameters of the model, are described such as in expression (1).

**Fig 2.**
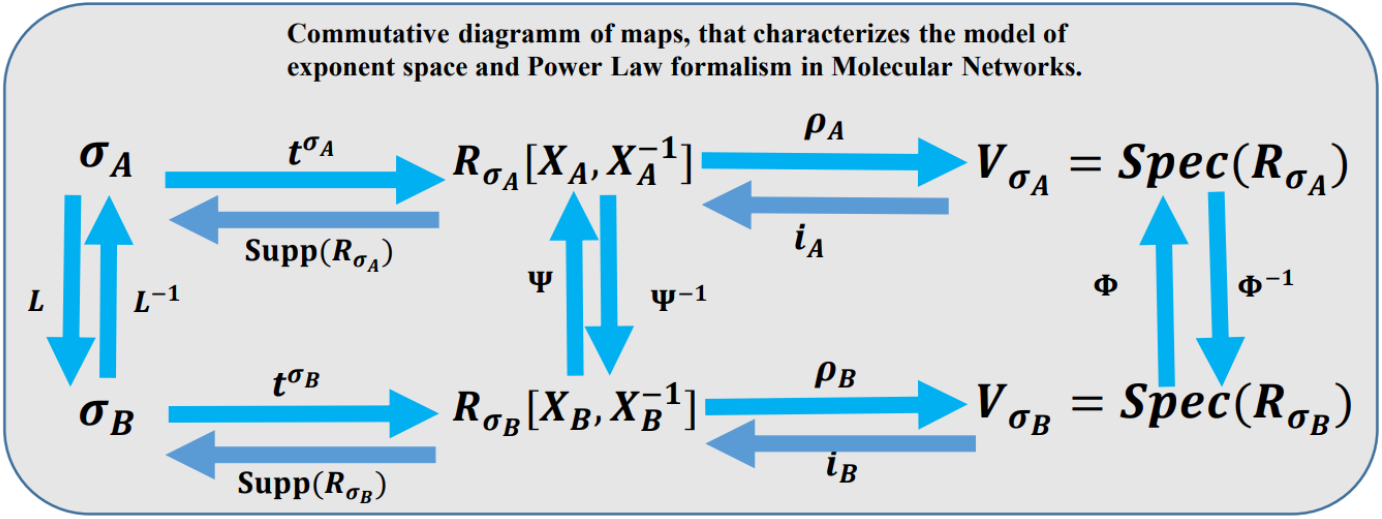
Commutative diagram of mappings between the exponent space, the ring of power-law polynomials for molecular networks, and the prediction of the number of binding sites. In formal mathematics, a diagram of maps is commutative if the composition of maps is consistent across all pathways. Each square in the diagram must commute, as explicitly we can see: 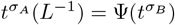, and,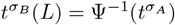. Similarly, the second square must also commute: *i*_*A*_(Ψ) = Φ(*i*_*B*_), and *i*_*B*_(Ψ^*−*1^) = Φ^*−*1^(*i*_*A*_). The definitions for, 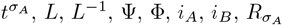, and 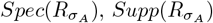 are provided in the Methods section. We assert that this diagram is commutative; see proof in, [34, 37] and supporting information.

At this point in our exposition, we will mention important theorems from the field of Algebraic Statistics and Combinatorial Algebraic Geometry.

### Relevant Theorems used in our methodology

Key results from Algebraic Statistics and Combinatorial Algebraic Geometry are essential here:

#### Birkhoff’s Theorem

Every doubly stochastic matrix is a convex combination of permutation matrices, [42].

#### Theorem (Algebraic Geometry)

Every magic square (a matrix whose rows and columns sum to the same quantity) is the convex hull of permutation matrices, which consist of their Hilbert bases, see [35].

### 0.1 1. Example: End-Product Inhibition Mechanism

In this study, we analyze the genetic regulation mechanism of end product inhibition, commonly found in bacteria and humans (see https://biocyc.org/pathway?orgid=ECOLIid=P4-PW, write in the prompt: aspartate superpathway). The differential equations governing the dynamics of *n* components under the power-law formalism are provided in [26].

For instance, when the pathway length is *n* = 3, there is one intermediate. For *n* = 9, there are seven intermediates. Examples include aspartate regulation by lysine, with 7 intermediates and 8 binding sites in the regulatory sequence structure, and other cases such as homoserine regulation by methionine with 3 intermediates and 3 binding sites. These mechanisms are highly relevant in amino acid biosynthesis (see https://biocyc.org/pathway?orgid=ECOLIid=P4-PWY (write in the prompt: aspartate superpathway) and https://www.britannica.com/science/metabolism/End-product-inhibition).

## Results

### Using Monoids in the Power Law Modeling

Using Monoids in the Power Law Formalism Based on the mathematical modeling of the power-law for genetic-metabolic networks and combinatorial algebraic geometry, we studied the geometric properties of the exponent space in the polynomials of the power-law modeling and their biological implications. The exponent space is associated with two aspects: first, in the context of genetic networks, it corresponds to the number of binding sites in the loci of the active sites in a transcriptional unity; second, in the context of metabolic networks, it corresponds to reaction orders. This analysis leads to extended applications in computational biology. Below, we present a mathematical formalism that supports our methodology (see Figure 2).

The workflow diagram in Figure 3 illustrates the computational methodology used to predict binding sites.

**Fig 3.**
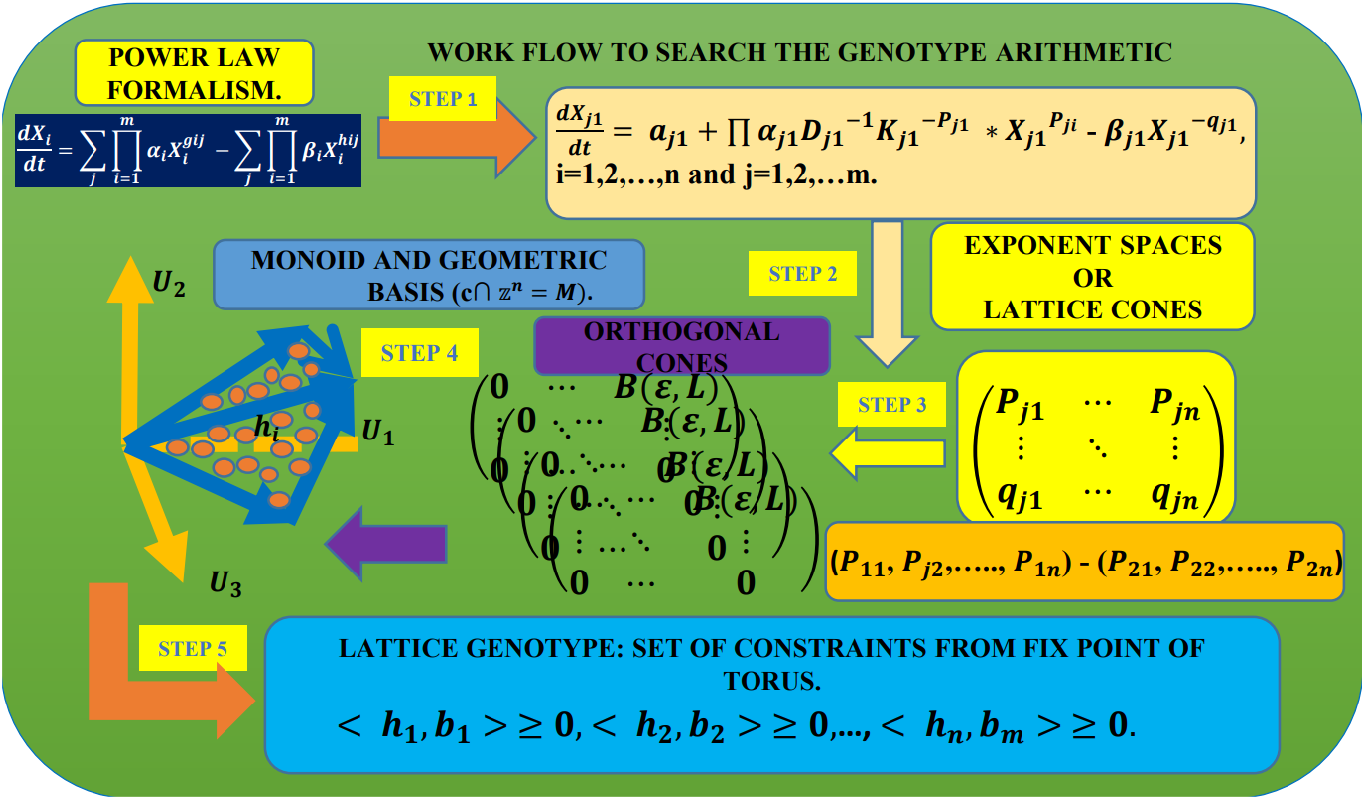
Workflow using Monoids in the Power Law Modeling. **Step 1:** Define the rate laws in the Power Law Modeling using the general mass-action expressions given in Equation (1) (Methods and Models section). **Step 2:** Construct matrices from the exponent space of nonlinear polynomials. Rows are ordered lexicographically [35, 36, 39]. Pairwise subtractions of the rows are performed to build a convex geometry [37]. These matrices are referred to as lattice-cones.**Step 3:** Compute orthogonal matrices from the lattice-cone matrices using random number generators based on the Boltzmann distribution [45–53]. **Step 4:** Calculate geometric bases using the Normaliz platform [44]. These geometric bases define a Monoid. **Step 5:** Use formulas (3) to compute fixed points on the torus [34]. These fixed points represent a family of constraints that define bounds for the number of binding sites.

### Using Monoids to approach a Family of Constraints (Arithmetic Genotype) for Binding Sites

Our methodology uses the workflow in Figure 3, supported by combinatorial algebraic geometry algorithms via the Normaliz software (https://www.normaliz.uni-osnabrueck.de/) [44]. Two theorems (detailed in the Methods section) are central to our application. Specifically, they demonstrate that the geometric bases for kinetic orders in the case of end-product inhibition are equivalent to permutation matrices over the identity matrix of dimension n×n, where n corresponds to the length of the metabolic pathway in a regulatory cascade.

Using the Boltzmann distribution [45–53], we systematically choose the best interaction. This geometric Monoid-based technique enables the construction of genetic interaction matrices. These matrices contain entries that represent the genotype-phenotype functional ratio:

Genotype/Phenotype == #*BS/*#*X*_*i*_ == Number of Binding Sites / Number of molecules in the network == Probability of binding sites occupied by the TF represented by the *X*_*i*_.

This ratio represents the probability of a transcription factor (TF) interacting with DNA. If this probability is maximal compared to other ratios in the row of the interaction matrix, it corresponds to the peak molar concentration of the phenotype *X*_*i*_.

With the exponents in the expression of the limit formula (3), a new space of constraints was computed, they define a lattice structure (an ismophormic to the n-dimensional integer space, Figure 2). In this point, we explained the symbol;

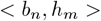

in the context of molecular biology. In mathematics is defined as the inner scalar product, as is shown below, The family of the found constraints in the step 5, let us to find numerical values for the number of binding sites *b*_*k*_’s, once what is computed the geometrical basis *h*_*k*_’s (Normaliz), the following set of inequalities represent a system of algebraic equations, see expression (3): where the geometrical bases computed from Normaliz software are called the Hilbert basis of the monoid, see [37, 39].

### Application of genotype arithmetic

For regulatory cascade lengths (e.g., n=7): Predicting binding sites in end-product inhibition mechanisms

The interaction between molecules participating in an allosteric mechanism of end-product inhibition has been modeled (Fig. 4). For cases ranging from n=3 to n=7, M.A. Savageau demonstrated that molecular phenotypes exhibit stable dynamics. However, for metabolic pathways longer than n=9, molecular products are not sustained (see [35]) and decay into unstable concentrations. In other words, for pathways of this length, the enzymatic kinetics of these products are inefficient due to energy conservation (e.g., ATP, ADP) and the optimal distribution of the cell’s resources. Additionally, a longer pathway results in delayed reaction times in response to biochemical signaling caused by environmental or exogenous biochemical changes.

**Fig 4.**
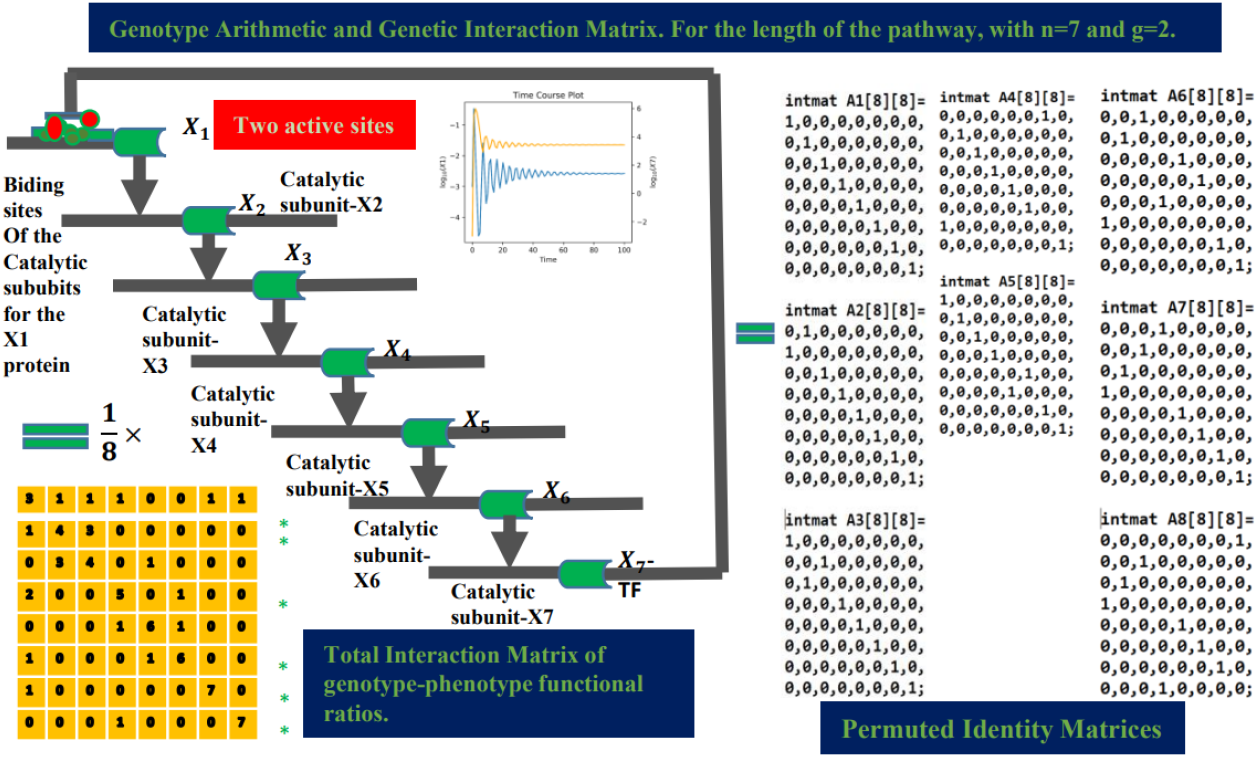
Case of end-product inhibition for the pathway length n = 7 (lysine-aspartate) with two active sites (red dots), see (sites cysteine 135, histidine 275 loci), https://www.uniprot.org/uniprotkb/P0A9Q9/entry, https://biocyc.org/pathway?orgid=ECOLIid=PWY0-781 (write in the prompt: aspartate superpathway), and https://www.britannica.com/science/metabolism/End-product-inhibition.

A detailed analysis of the regulatory cascade for n=7 is presented below, while cases for n=3 to n=9 are provided in the supplementary material. The workflow for the n=7 case is described as follows. Figure 4 shows the cascade network for the reactions involved in the regulation of aspartate by lysine, regulated by two active sites, red dots in figure 4 (see: sites cysteine 135, histidine 275 loci) https://www.uniprot.org/uniprotkb/P0A9Q9/entry and https://biocyc.org/pathway?orgid=ECOLIid=P4-PW (write in the prompt: aspartate superpathway)). Here, the intermediate interactions are catalytic reactions (metabolism), while the end product, *X*_7_ antagonizes the activity of the allosteric protein *X*_1_ (gene regulation).

The respective geometric bases of the exponent space associated with each transcription factor *X*_*i*_ were calculated, along with an additional matrix for the constraint D. By summing these matrices (the convex hull of the network), a bistochastic matrix was obtained. Dividing the rows by 8 (7 molecules on network plus 1 constraint) ensured that each row summed to 1. We also present the monoid and lattice generated for this case, as defined by formula (3).

The graphs correspond to the molar concentration rates for the initial product *X*_1_ (blue) and the final product *X*_7_ (yellow), where *X*_7_ antagonizes the activity of the allosteric protein *X*_1_. The yellow matrix weights each entry with the functional genotype/phenotype ratio, defined as: genotype/phenotype ratio = binding sites/phenotypes. Rows with ratios that reproduce the binding site g for the allosteric protein *X*_1_ are marked with an asterisk.

The blue matrix in the figure above corresponds to the total sum of all monoid generators, as shown in Figures 5 and 6 (fourth column). This matrix is divided by the total number of quantitative phenotypes: nine molecules in the network plus one constraint, for a total of 8. The entries of the blue matrix represent the genetic functional rate.

**Fig 5.**
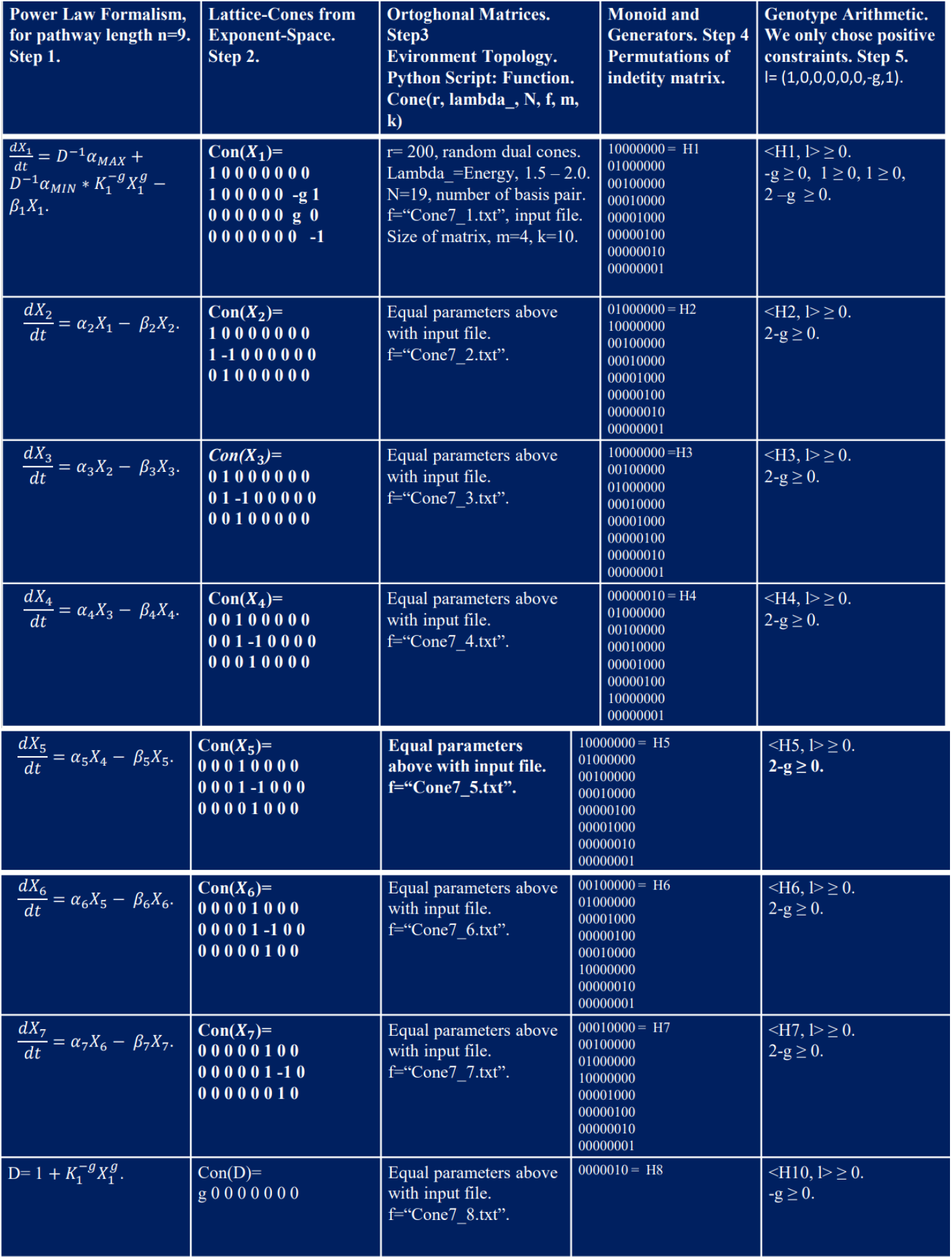
Analysis of the exponent space for the equations *X*_1_ up to *X*_9_ applied to the case length of the pathway n=7. * Each column refers to steps into the workflow diagram in figure 3.

**Fig 6.**
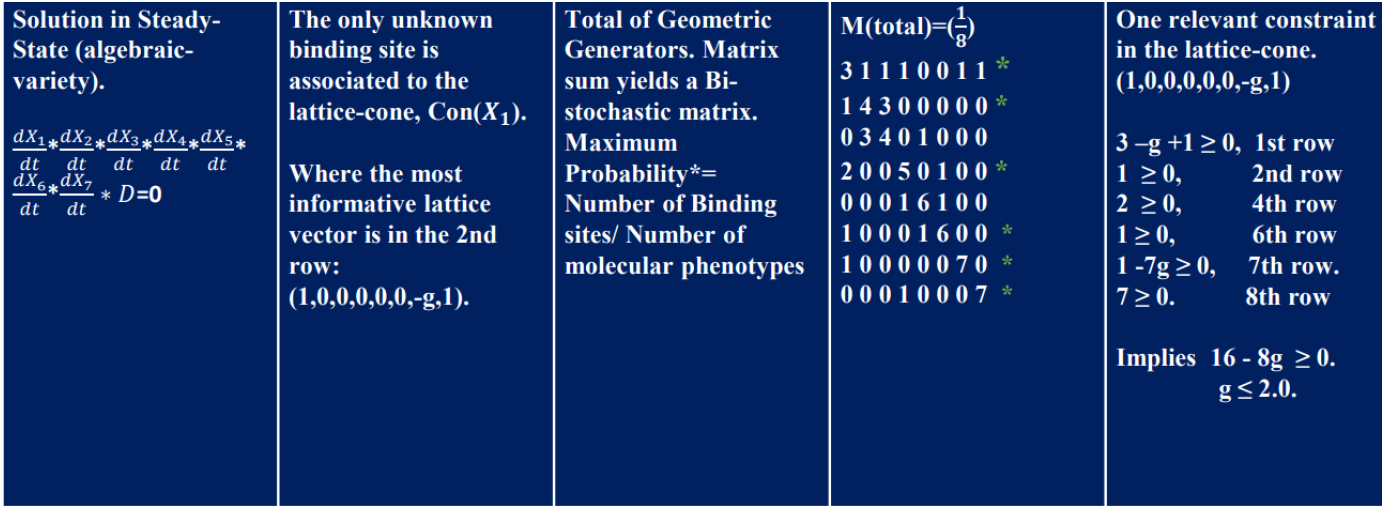
Analysis of the exponent space for constraint equation D.

This corresponds to a bistochastic matrix, meaning that each entry represents the probability of interaction between elements in the network. Notably, the sum of elements for each row and column equals 1.

In the following tables, each column corresponds to a step in the pipeline previously defined for the n=7 case.

The first columns in Figures 5 and 6 describe the full-system equation for the pathway length *n* = 7, where only the first equation associated with *X*_1_ contains the unknown binding site g. In column 2, we construct the lattice-cones with rows corresponding to each exponent (lattice-vector) for each monomial term in the equation. The intermediate rows represent the exponents of the monomials when the equations are factored, as shown in the first column.

In the third column, we simulate orthogonal random matrices for the lattice-cones using the Python function *Cone*(*r*, *N, f, m, k*) (see supporting information), where *f* refers to text files containing the matrices built in column 2. These files serve as input data for computing geometric generators. The generators represent a Monoid structure and are computed using the Normaliz platform (see [44]). The output files are identity permutation matrices, which are displayed in the fourth column.

Finally, using the inequalities set in expression (3), we generate the arithmetic genotype for each local geometry, as shown in the fifth column in Figures 5 and 6.

In Figure 6 (first column and last row), the product of the full-system equations in steady-state is shown. In algebraic geometry, this corresponds to solving algebraic non-linear polynomials. According to the monomial map, these equations yield a set of lattice-cones. The most informative of these corresponds to *Con*(*X*_1_), whose lattice vector (1, 0, 0, 0, 0, 0, *g*, 1) contains the unknown binding site g.

To predict its value, we sum the complete set of matrices for each Monoid in the fourth column, yielding the total matrix M. Dividing M by the number of molecular phenotypes in the network (8 in this case), plus one constraint equation D, results in a bistochastic matrix.

Rows 1, 2, 4, 6, 7, and 8 produce the maximum projection (scalar inner product or maximum peak of molar concentration) with the lattice vector (1, 0, 0, 0, 0, 0, *g*, 1), consistent with formula (3). Thus, we derive an arithmetic representation of the genotype for the entire network. By summing the complete set of inequalities, we obtain a range for *g*.

### Arithmetic Genotype using monoids reduces computational complexity versus Power Law Modeling

The values for the number of binding sites *g* associated with each pathway length in Figure 7 correspond to the only stable physiologies for end-product inhibition, as reported previously in [26]. The molar concentration curves were plotted using the DST3 software [54].

**Fig 7.**
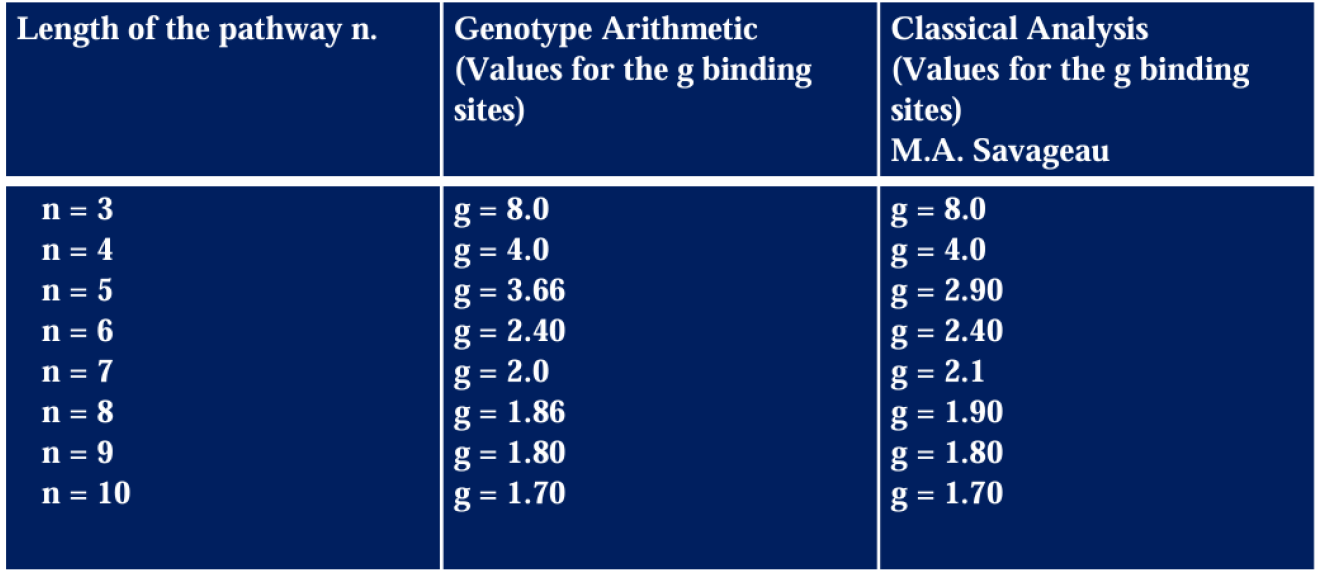
Comparison of the predicted values for the classic analysis of power low formalism and genotype arithmetic using monoids. * In the case of rational values of *g*, all integer monoid can be extended to a rational monoid, see [37].

We note that the results of our work elucidate a deep connection between the exponent space of the number of binding sites and the quantitative phenotypes of molar concentration variables *X*_*i*_. In the case of this allosteric network, the generators are represented by a permuted identity matrix. At this point, it is important to mention that the construction of this matrix was performed manually.

We observe the approximation for the *g* values for the cases *n* = 5 and *n* = 7. The difference in values here may be attributed to the construction of the permuted identity matrices. It is possible, that more precise permutations of the identity matrix are required. With this refinement, we could predict precise values for other pathway lengths.

The Normaliz program computed only sections of the identity matrix in the output files, as shown in the supplementary material. In each iteration, it computes only the Hilbert basis corresponding to a face of the lattice cone or polytope lattice. To construct the complete permuted identity matrix, we justify our construction for all faces using the theorems mentioned in the final part of the Methods and Models section. For instance, for the case *n* = 9, we describe the following output file from Normaliz in the supplementary material.

The family of matrices governing the exponent space corresponds to the symmetric group of the identity matrix in the case of end-product inhibition. It describes the hidden geometry in the lattice vectors formed by the number of binding sites *g*. For different cases of genetic and metabolic networks, exploring the family of combinatorial matrices that govern genotype arithmetic will be an area of interest for future work.

In the following, we present the growth curves for the first allosteric enzyme *X*_1_ and the end products *X*_3_, *X*_4_, *X*_5_, *X*_6_, *X*_7_, *X*_8_ and *X*_9_. We simulated the plots for all stable cases of pathway lengths, and the plots are shown below, see Fig. 8.

**Fig 8.**
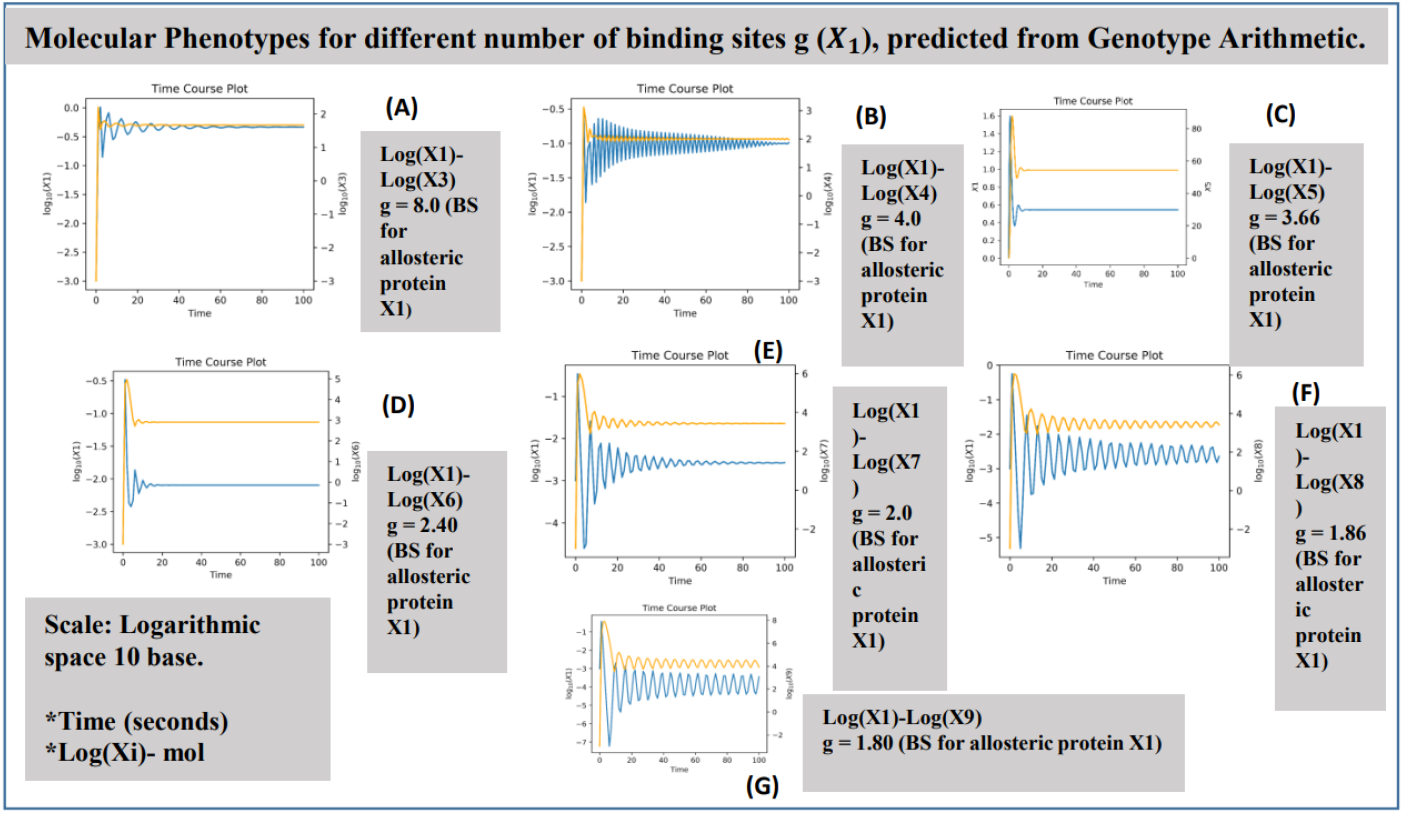
Time course plot for length of the pathways. A) n=3, B) n=4, C) n=5, D) n=6, E)n=7, F) n=8, G) n=9. Design Space Toolbox V. 3.0. was used for the plots shown above, see [54]. The plots quantify the molar concentration for the production of the end product (yellow) and the inhibition of the first allosteric protein in the metabolic cascade (blue). In all cases, the flow of biomass is sustained as stable quantitative phenotypes, as predicted in [26]. The stability is governed by the predicted value for the binding sites *g*, shown in Table 3, where we compare all the predicted values for both methodologies. The plots also show that the concentration rate for the end product increases over time before decreasing to repress the allosteric protein. Simultaneously, the concentration of the first protein decreases due to antagonism by the action of the end-product enzyme *X*_*n*_ with *n* = 3, 4, 5, 6, 7, 8, 9.

In this final part of the section, we highlight the significant potential for reducing computational complexity when solving kinetic orders in large networks. This reduction is based on the fact that the geometric local basis (Hilbert basis) has been shown to converge in polynomial time [40].

Each molecule in the network has a local matrix basis, and its computation belongs to the polynomial time class. By solving the network in local modules (varieties) and dividing it into monoids, it becomes possible to solve each module in polynomial time. This modular approach significantly reduces the computational complexity required to determine the full set of kinetic orders.

## Discussion

Mathematical and computational formalisms are central to molecular network modeling, requiring robustness to enable the large-scale experiments currently underway. In future work, the Monoid technique could be extended to algebraic varieties (see [34–38]), as shown in Figure 2. This extension could model large blocks of biochemical systems using local varieties, such as artificial modules, to solve large numbers of parameters within the network.

In Figure 2, it is also demonstrated that the Power Law Modeling has a deep connection to algebraic varieties, specifically toric varieties. This connection enabled the analysis and prediction of the number of binding sites using fixed points on the torus (see [34]).

The main result of this work is related to the properties of the exponent space derived from the Power Law Modeling for biochemical networks. These properties may provide a characterization of the relationship between **genotype and phenotype** through the monomial map 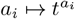 (see [34, 37]). Specifically, the *a*_*i*_’s correspond to the number of binding sites associated with the genomic loci of functional sites in regulatory genome sequences, while the monomials 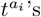 represent kinetic variables of molar concentrations of molecular phenotypes (see [26–32]).

Additionally, the fixed points in the exponent space, expressed in Equation (3), define a lattice structure. This lattice is related to an arithmetic of constraints on the number of binding sites in DNA interactions or the regulatory subunits of allosteric proteins, [41]. The Monoid structure defining these constraints is specific to each case of molecular interaction and may define an **Arithmetic of the Genotype** for these families of monoids related to molecular networks.

The Monoid technique in molecular networks also defines combinatorial operations on the lattice (see Figure 2), which is formed by the number of binding sites. These combinatorial operations such as subtractions, additions, scalar inner products, and lexicographic ordering among lattice vectors (see [34–38]) can reflect and characterize genomic and biotechnological techniques, including gene insertions, deletions, point mutagenesis, transpositions, and transposon libraries. Moreover, the fixed points in the exponent space may characterize different families of lattice vectors (see [34–38]), representing binding sites associated with various configurations of regulatory genome sequences. This suggests that monoids could reveal new combinatorial properties of regulatory sequences and their interactions with the biochemical environment.

The combination of both methodologies Power Law Modeling and Arithmetic of the Genotype is applicable to molecular interactions, such as transcription factor-DNA interactions and protein-protein interactions (see [**?**, 45–49, 51–53]), and could eventually be extended to large-scale molecular networks. The scheme in Figure 2 also provides a rigorous tool to solve the space of kinetic orders in molecular networks, piece by piece, using local varieties. By associating a Monoid geometry with each molecule in the network and then combining these local geometries into a complete algebraic variety (see [34–38]), this approach could significantly reduce computational complexity by solving parameters through local varieties or geometric modules.

Finally, a future application of this allosteric mechanism could be explored. Previously, a gene and metabolic pathway associated with the TGF-*β* system *in Human* was outlined. A regulatory cascade is activated by end-product inhibition in the TGF-*βI* receptor protein. This regulatory pathway is relevant to idiopathic pulmonary fibrosis (see [55]).

## Conclusion

This methodology revises combinatorial properties related to genetic and metabolic networks using the Power Law Modeling. One of these properties is the technique of Monoids. Using this algebraic and geometric object, a Genotype Arithmetic was constructed, related to a set of constraints that define a range of values for predicting the number of binding sites in an allosteric enzyme. This technique was applied to the allosteric system of End-Product Inhibition. The predicted values were similar to those previously reported in [26].

The utility of the Monoid technique opens up the possibility of analyzing large genetic and molecular networks in the future. Its connection with algebraic varieties could model a large space of kinetic orders by studying each molecule in the network as a module governed by local varieties or local Monoids.

## Supporting information

https://github.com/mpolocv/End-Product-Inhbition (see Supplementary Materialv1.txt) and https://mybinder.org/v2/gh/Normaliz/NormalizJupyter/master

## Acknowledgments

Marco Polo Castillo Villalba is a doctoral student from the Programa de Doctorado en Ciencias Biomédicas, Universidad Nacional Autónoma de México (UNAM) and has received CONAHCYT fellowship 754050. We acknowledge funding from UNAM and from the National Institutes of Health (grant number 5R01GM110597-03), and funding support from INER. Also, we give our thanks for all your advice and comments Luis Mendoza, and Yalbi I. Balderas Martínez, specially a huge thanks to Michael A. Savageau, for all the discussions and technical suggestions, during my scholar stay with him to Julio Collado-Vides, Osbaldo Resendis-Antonio and Pedro Miramontes-Vidal.

